# Corilagin attenuates high glucose-induced neurotoxicity and mitochondrial dysfunction through restoration of the AMPK-SIRT1-PGC1α-TFAM signalling axis

**DOI:** 10.64898/2026.07.26.740444

**Authors:** Rudra Chakravarti, Dipanjan Roy, Jhanshi Chigilipalli, Bireswar Bhattacharya, Mansi Arya, Manoj Manna, Dipanjan Ghosh

## Abstract

Mitochondrial dysfunction and oxidative stress represent two interconnected, primary causes for Diabetic Neuropathy (DN); however, the majority of currently available anti-diabetic therapies have focused on glucose control as opposed to neurodegenerative downstream effects. Corilagin, is an ellagitannin having high anti-oxidant properties; however, it has not been evaluated against hyperglycemia induced neuronal injury. The present study demonstrates the ability of Corilagin to protect against mitochondrial dysfunction via models of diabetic nephropathy and cerebral ischemia. High glucose (50 mM, 24 hr) was utilized to induce diabetes like conditions in the SH-SY5Y human neuroblastoma Cell Line. High glucose induced significant decreases in cell viability, increases in intracellular and mitochondrial reactive oxygen species, depletion of reduced glutathione reserves, induces apoptosis, and causes mitochondrial depolarization and fragmentation. Corilagin pre-treatment attenuated each of these high-glucose induced effects by protecting against mitochondrial membrane potential loss and maintaining mitochondrial network morphology while reducing apoptotic cell fraction relative to glucose alone. Additionally, these protective effects were accompanied by restoration of AMPK phosphorylation and up-regulation of SIRT1, PGC1α and TFAM, components that are part of the principal signaling pathway that regulates mitochondrial biogenesis; therefore, therefore, this pathway may contribute mechanistically to the cyto-protective effect of Corilagin.

**Graphical abstract.**
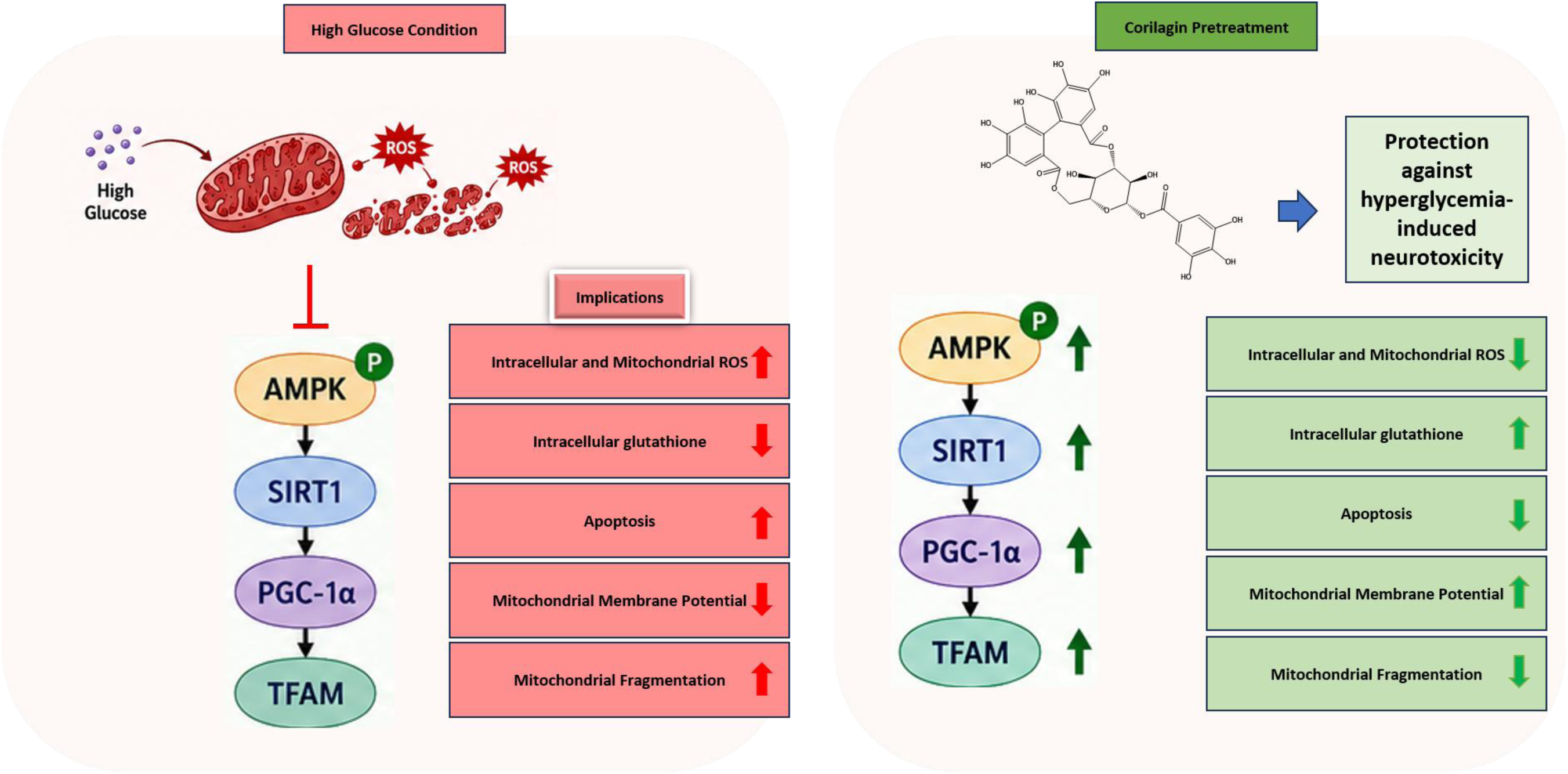
Proposed mechanism underlying the neuroprotective effects of Corilagin against high glucose-induced mitochondrial dysfunction. High glucose suppresses AMPK phosphorylation, leading to downregulation of the SIRT1–PGC-1α–TFAM signaling axis, increased intracellular and mitochondrial reactive oxygen species (ROS), glutathione depletion, apoptosis, mitochondrial depolarization, and mitochondrial fragmentation. Corilagin pretreatment restores AMPK activation and the downstream SIRT1–PGC-1α–TFAM pathway, thereby reducing oxidative stress, preserving intracellular glutathione, preventing apoptosis, maintaining mitochondrial membrane potential, and protecting mitochondrial network integrity.

## 1. Introduction

Diabetes Mellitus is now one of the world’s major public health problems. The International Diabetes Federation estimates that 589 million adults will live with diabetes in 2024, and that number is expected to increase to 853 million by 2050 [1]. Diabetic Neuropathy (DN), or peripheral neuropathy caused by long-term exposure to elevated levels of glucose that damages the nerves and vascular walls, affects up to half of all individuals who suffer from diabetes for at least 25 years, and it is currently the most common cause of non-traumatic lower limb amputations throughout the world [2].

The exact etiology of DN is quite complex; however, increasing evidence supports the idea that hyperglycemia induced oxidative stress is the primary unifying process involved in the pathogenesis of DN [3]. In support of this concept, Brownlee’s "unifying hypothesis" states that excessive intracellular glucose overwhelms the mitochondria electron transport chain causing the production of vast amounts of superoxides [4]. This mitochondrial dysfunction causes a cascade of additional damage including the formation of Advanced Glycation End Products (AGEs), activation of the polyol pathway, and reduction of endogenous antioxidant defenses. Due to their inability to modulate glucose uptake, neurons are especially vulnerable to the effects of glucotoxicity, apoptosis, and loss of function [5]. As such, therapeutic agents capable of neutralizing mitochondrial reactive oxygen species (ROS) and restoring metabolic balance appear to be a promising approach to preserving neuronal integrity in diabetes.

The current treatments of DN are primarily based on controlling blood sugar levels and managing symptomatic pain. Current treatment options do nothing to halt the continued degeneration of neurons [6, 7]. Historically, antidiabetic drugs were developed to protect organs critical to metabolism such as the liver, pancreas, and kidneys. As compared to neuronal and cardiovascular targets little emphasis was placed on developing drugs that could mitigate the oxidative and mitochondrial processes involved in complications related to diabetes such as DN [8].

Additionally, many conventional treatments used to treat DN can be challenging to use due to severe side effects [9, 10]. Therefore, several categories of naturally occurring compounds such as alkaloids, flavonoids, and tannins have demonstrated improved glucose uptake and insulin sensitivity while also demonstrating inhibition of oxidative stress and activation of AMPK signaling [11–14]. As such, naturally derived compounds appear to be an appealing option for treating multiple aspects of diabetic complications such as DN.

There are several polyphenols and flavonoids that have demonstrated efficacy in preclinical models of DN. Luteolin has been demonstrated to improve nerve conduction velocity while decreasing markers of oxidative stress in diabetic rats [15]. Rutin has been shown to decrease plasma glucose concentrations and oxidative stress through the Nrf2 pathway in diabetic rat models [16]. Resveratrol and pinocembrin have each demonstrated enhancement of mitochondrial function while reducing oxidative damage in diabetic animal models [17, 18].

Corilagin (β-1-O-galloyl-3,6-(R)-hexahydroxydiphenoyl-D-glucose), an ellagitannin found in medicinal plants such as Phyllanthus niruri and Terminalia chebula, has also been identified as a potentially viable candidate for preventing DN. Corilagin has been shown to exhibit significant antioxidant, anti-inflammatory and hepatoprotective properties [19]. Studies have recently demonstrated the utility of Corilagin for the prevention of diabetic nephropathy. Specifically, Corilagin was demonstrated to prevent podocyte injury in diabetic nephropathy by activating the SIRT1-AMPK signaling axis, which plays a key role in regulating cellular energy homeostasis and autophagy [20], and Corilagin has been shown to exert neuroprotective effects in cerebral ischemia by reducing oxidative stress via the Nrf2 pathway [21].

Although these results demonstrate promise for kidney disease and ischemic brain models, there is no information available regarding the ability of Corilagin to protect neurons from the acute oxidative burst generated by hyperglycemia in diabetic neuropathy. In this study, we examined the cytoprotective potential of Corilagin using human SH-SY5Y neuroblastoma cells as an in vitro model of glucotoxic neuronal injury. We predicted that Corilagin would act as a metabolic rescuer by alleviating high glucose-induced apoptosis through elimination of intracellular ROS, maintenance of mitochondrial function, and reactivation of the AMPK-SIRT1-PGC1α-TFAM signaling axis.

## 2. Experimental materials and methods

### 2.1 Materials

Dulbecco’s Modified Eagle Medium (DMEM, low glucose, 1 g/L; Gibco, Cat. No. 31600034), Fetal Bovine Serum (FBS; Gibco, Cat. No. 10270106), Antibiotic-Antimycotic (100×; Gibco, Cat. No. 15240062), Dulbecco’s Phosphate Buffered Saline (DPBS; Gibco, Cat. No. 14190144), Trypsin-EDTA (0.25%), phenol red (Gibco, Cat No.25200056), D-(+)-Glucose (Sigma-Aldrich, Cat. No. G7021), D-Mannitol (Sigma-Aldrich, Cat. No. M1902), Corilagin (Toronto Research Chemicals, Cat. No. C695400), MTT (Sigma Aldrich, Cat. No. 475989), 2′,7′-Dichlorodihydrofluorescein diacetate (HßDCFDA; Invitrogen, Cat. No. D399), MitoSOX™ Red Mitochondrial Superoxide Indicator (Invitrogen, Cat. No. M36008), ThiolTracker™ Violet (Invitrogen, Cat. No. T10095), Annexin V-FITC Apoptosis Detection Kit (Invitrogen, Cat. No. BMS500FI-100), JC-1 Mitochondrial Membrane Potential Probe (Invitrogen, Cat. No. T3168), and MitoTracker™ Deep Red FM (Invitrogen, Cat. No. M46753) were used in this study. SH-SY5Y human neuroblastoma cells were used as the in vitro model. It was procured from NCCS, Pune. Flow cytometric analyses were performed using a BD LSRFortessa™ flow cytometer (BD Biosciences, USA), and all the analysis were done using FlowJo-V10; while fluorescence imaging was carried out using a Leica Stellaris confocal laser scanning microscope (Leica Microsystems, Germany). RIPA buffer (Sigma-Aldrich, Cat. No. R0278). Primary antibodies against AMPKα (62 kDa; Rabbit recombinant monoclonal, ABclonal, Cat. No. A27740), phospho-AMPK (T182+T172) (62 kDa; Rabbit recombinant monoclonal, ABclonal, Cat. No. AP1441), SIRT1 (110 kDa; Rabbit recombinant monoclonal, ABclonal, Cat. No. A19667), TFAM (26 kDa; Rabbit recombinant monoclonal, ABclonal, Cat. No. A3173) and PGC1α (110 kDa; Rabbit recombinant monoclonal, ABclonal, Cat. No. A20995) were used for immunoblotting.

### 2.2 Cell culture

SH-SY5Y human neuroblastoma cells were cultured in Dulbecco’s Modified Eagle Medium (DMEM) containing low glucose (1 g/L). The culture medium was supplemented with 10% FBS and 1x Antibiotic-Antimycotic solution to maintain optimal growth conditions. Cells were maintained in a humidified incubator at 37 °C and 5% CO_2_.

### 2.3 Cell viability assay

Cell viability was assessed using the MTT assay [22] to evaluate the cytoprotective effect of Corilagin against high glucose-induced cytotoxicity. Briefly, SH-SY5Y cells were seeded in flat-bottom 96-well plates at a density of 1 × 10ß cells/well and allowed to adhere overnight. Initially, the cytotoxicity of Corilagin (2.5, 5, 10, 20, 50, and 100 μM) and glucose (5–250 mM) was evaluated individually following 24 h of treatment to determine their effects on cell viability. Based on these preliminary experiments, 10 μM Corilagin and 50 mM glucose were selected for subsequent cytoprotection studies. For these experiments, SH-SY5Y cells were seeded in 96-well plates at a density of 1 × 10ß cells/well and allowed to attach overnight. On the following day, the culture medium was replaced with fresh serum- and antibiotic-free medium. Cells were pretreated with 10 μM Corilagin for 1 h, followed by exposure to 50 mM glucose for the indicated duration. Equimolar concentrations of mannitol were included as osmotic controls in the respective experimental groups. Following treatment, 10 μL of MTT solution (5 mg/mL) was added to each well, and the plates were incubated at 37°C for 4 h. The plates were then centrifuged, the supernatant was carefully discarded, and the resulting formazan crystals were dissolved in 200 μL dimethyl sulfoxide (DMSO). Absorbance was measured at 570 nm using a microplate reader, and cell viability was expressed as a percentage relative to the untreated control.

### 2.4 Measurement of Cellular and mitochondrial ROS

Intracellular reactive oxygen species (ROS) levels induced by glucose exposure were quantified utilizing H2DCFDA staining [23]. Within the cytoplasm, esterases cleave the dye, and subsequent oxidation by ROS converts it to the highly fluorescent 2’,7’-dichlorofluorescein (DCF) (indication of ROS generation). Briefly, SH-SY5Y cells were plated in 12 well plates at a density of 1x10^5^ Cells/well in the growth media as discussed previously. After 24 hours incubation, the culture media was discarded and 1ml of fresh media (without Serum and antibiotic-antimycotics) was added to each well. Cells were then pretreated with Corilagin at a final concentration of 10 µM for 1 hour prior to exposure to high glucose conditions. Glucose solution was then added to a final concentration of 50 mM. Equimolar concentration of mannitol was administered as an osmotic control across the experimental groups. Following the respective experimental treatments, cells were harvested by trypsinization, washed with 1× DPBS, and incubated with 20 μM HßDCFDA at 37°C for 1 h. Following incubation, cells were washed once with 1× DPBS to remove excess dye and resuspended in 1× DPBS for analysis. Intracellular ROS levels were quantified using a BD LSRFortessa™ flow cytometer. HßDCFDA fluorescence was detected using the FITC channel, and a minimum of 10,000 events were acquired for each sample. Data were expressed as mean fluorescence intensity (MFI), representing intracellular ROS levels. Mitochondrial-specific oxidative stress was assessed using MitoSOX, a fluorescent indicator targeting mitochondrial superoxide. Stressed mitochondria with high superoxide levels were identified by the accumulation of the dye and the generation of Fluorescence. Cells were treated the same way as the cellular ROS experiment. Following the respective experimental treatments, cells were harvested by trypsinization and incubated with 5 μM MitoSOX™ Red at 37°C for 1 h. The cells were then washed twice with 1× DPBS to remove excess dye and resuspended in 1× DPBS for flow cytometric analysis. Mitochondrial superoxide levels were quantified using a BD LSRFortessa™ flow cytometer. MitoSOX™ Red fluorescence was and detected in the PE channel. A minimum of 10,000 events were acquired per sample, and the MFI was used as an indicator of mitochondrial superoxide generation.

### 2.5 Reduced glutathione assay

To evaluate the protective effects of Corilagin against depletion of intracellular antioxidant reserves, cellular reduced thiols (primarily glutathione, GSH) were quantified using ThiolTracker™ dye. SH-SY5Y cells were seeded in 12-well plates at a density of 5 × 10ß cells/well and treated with Corilagin (10 μM) and high glucose (50 mM) as described above. Following the respective experimental treatments, cells were harvested by trypsinization, washed once with 1× DPBS, and stained with ThiolTracker™ according to the manufacturer’s instructions to assess intracellular reduced glutathione (GSH) levels. After incubation, cells were washed with PBS to remove excess dye and resuspended in 1× PBS for flow cytometric analysis. Fluorescence was acquired using a BD LSRFortessa™ flow cytometer in Qdot 525 detection channel. 10,000 events were acquired for each sample, and the MFI was used as an indicator of intracellular GSH levels.

### 2.6 Annexin V- FITC assay for assessing Apoptosis

The protective effect of Corilagin against high glucose-induced apoptosis was evaluated using a flow cytometry-based Annexin V assay. Briefly, SH-SY5Y cells were seeded in 12-well plates at a density of 1 × 10ß cells/well in complete growth medium and allowed to adhere overnight. The following day, the culture medium was replaced with 1 mL of serum- and antibiotic-free medium. Cells were pretreated with 10 μM Corilagin for 1 h, followed by exposure to 50 mM glucose for 24 h. Equimolar mannitol was used as an osmotic control where appropriate. Following treatment, cells were harvested by trypsinization, washed twice with cold 1× PBS, and stained with Annexin V and propidium iodide (PI) according to the manufacturer’s instructions. Data were analyzed using a bivariate dot plot, where FITC fluorescence (Annexin V-FITC) was displayed on the X-axis and PE-CF594 fluorescence (Propidium Iodide, PI) on the Y-axis. Quadrant gating was applied to distinguish viable cells (Annexin V−/PI−), early apoptotic cells (Annexin V+/PI−), late apoptotic cells (Annexin V+/PI+), and necrotic cells (Annexin V−/PI+), and the percentage of cells in each population was quantified.

### 2.7 Mitochondrial membrane potential and morphology analysis

Mitochondrial membrane potential (Δψ_m_) and polarity were evaluated employing JC-1 dye staining [24] following manufacturer’s protocol. In healthy cells with high membrane potential, the dye accumulates in the mitochondria and forms J-aggregates, emitting red fluorescence. Conversely, in unhealthy or apoptotic cells with diminished membrane potential, the dye disperses into the cytoplasm as monomers, emitting green fluorescence. SH-SY5Y cells were seeded in glass-bottom dishes at a density of 5 × 10³ cells/well and allowed to adhere overnight. Cells were pretreated with 10 μM Corilagin for 1 h, followed by exposure to 50 mM glucose for 24 h. Subsequently, the cells were incubated with 2.5 μM JC-1 dye for 30 min at 37°C, washed with 1× DPBS to remove excess dye, and immediately imaged using a confocal microscope. JC-1 fluorescence was recorded at 485 nm excitation and 515 nm emission, and mitochondrial membrane potential was evaluated based on changes in JC-1 fluorescence. Mitochondrial morphology was assessed using MitoTracker™ Deep Red staining. SH-SY5Y cells were seeded and treated as described for the JC-1 assay. Following the respective treatments, cells were washed with 1× PBS and incubated with MitoTracker™ Deep Red according to the manufacturer’s instructions for 30 min under standard culture conditions. The cells were then washed twice with fresh 1× DPBS to remove excess dye and immediately imaged using a confocal microscope. Fluorescence was acquired at an excitation wavelength of 644 nm and an emission wavelength of 665 nm. Quantitative image analysis was performed using LasX 2D analysis software to evaluate mitochondrial morphology by measuring mitochondrial area (μm²), roundness, aspect ratio (AR), and maximum Feret diameter (μm) across the different treatment groups.

### 2.8 Western blot analysis

To investigate the mechanistic activation of the AMPK-SIRT1-PGC1α signaling cascade, protein expression profiling was performed via Western blot analysis. SH-SY5Y cells were seeded in 60-mm culture dishes at a density of 1 × 10ß cells/dish and treated with Corilagin and glucose as described above. Following the respective treatments, cells were washed twice with ice-cold PBS and lysed in RIPA buffer (Sigma-Aldrich, Cat. No. R0278) supplemented with protease and phosphatase inhibitor cocktails. Total protein lysates were collected for subsequent Western blot analysis. Protein concentrations were quantified utilizing a BCA protein assay kit. Equal amounts of total protein from each sample were denatured by boiling in Laemmli loading buffer, resolved via SDS-PAGE on appropriate percentage gels (10%), and subsequently electrophoretically transferred onto polyvinylidene difluoride (PVDF) membranes. To prevent non-specific antibody binding, the membranes were blocked with 5% Bovine Serum Albumin (BSA) in Tris-buffered saline containing 0.1% Tween-20 (TBST) for 1 hour at room temperature. The blocked membranes were then washed and incubated overnight at 4°C with specific primary antibodies. Following rigorous washing with TBST, the membranes were incubated with the corresponding HRP-conjugated secondary antibodies for 1 hour at room temperature. Immunoreactive protein bands were visualized using an enhanced chemiluminescence (ECL) detection system and captured using a digital imaging system (Biorad ChemiDoc XRS+ System). β-actin (42 kDa) was utilized as the housekeeping gene. Densitometric quantification of the protein bands was performed to evaluate the relative expression levels of the target proteins.

### 2.9 Statistical significance

Statistical analysis and the generation of graphical data were performed to evaluate significance between treatments. Experiments were performed in triplicate if not mentioned otherwise. Comparative analyses between treatment groups (e.g., control, mannitol, glucose alone, and Corilagin pretreatment variations) were conducted to establish statistical significance using One-way ANOVA, which was denoted using standard asterisk notations corresponding to the respective p-values.

## 3. Results

### 3.1 Corilagin attenuates glucose-induced cytotoxicity in SH-SY5Y cells

To investigate the potential cytoprotective effects of Corilagin against hyperglycemic stress, SH-SY5Y cells were exposed to high glucose concentrations, and cell viability was evaluated. The concentration of 50 mM glucose was selected based on our dose-response experiments (Figure 1A) and previous studies demonstrating that this concentration consistently induces oxidative stress, mitochondrial dysfunction, and apoptosis in SH-SY5Y neuroblastoma cells without causing overwhelming cytotoxicity [25]. Corilagin singly at 10uM had minimal cytotoxicity (∼80% viability) (Figure 1B). Osmotic control using equimolar mannitol (50 mM) demonstrated minimal impact on viability, confirming that the observed cytotoxicity was specifically due to glucose-induced stress (Figure 1C). Pretreatment with Corilagin for 1 hour prior to glucose exposure effectively rescued the cells. Specifically, 10 µM Corilagin significantly improved cell survival against 50 mM glucose challenges at 24 hours (Figure 1C; p < 0.001). This concentration is also consistent with previous studies showing that Corilagin in the 5–20 µM range is well tolerated by SH-SY5Y neuronal cells, while effectively exerting neuroprotective and antioxidant effects [26].

**Figure 1.**
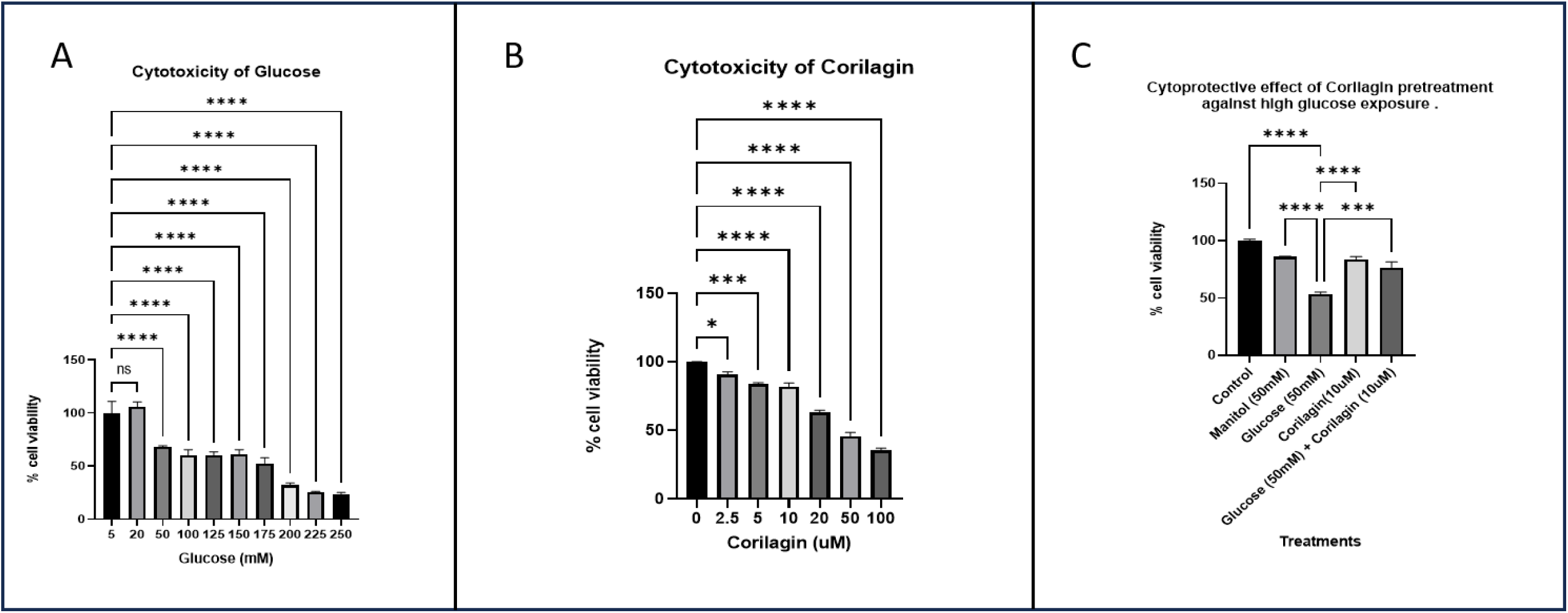
Dose-response evaluation and cytoprotective efficacy of Corilagin against high glucose-induced stress in SH-SY5Y cells. (A) Dose-dependent cytotoxicity of glucose. Cells were exposed to increasing concentrations of glucose (5 to 250 mM) for 24 hours, demonstrating a highly significant reduction in viability at 50 mM and above. (B) Dose-dependent evaluation of Corilagin toxicity. Cells were treated with Corilagin (0 to 100 µM) for 24 hours to establish dose for subsequent assays. (C) Cytoprotective effect of Corilagin pretreatment. Cells were pretreated with 10 µM or 20 µM Corilagin for 1 hour prior to a 24-hour challenge with 50 mM glucose. Mannitol (50 mM) was included as an osmotic control. Pretreatment with 10 µM Corilagin significantly preserved cell viability, rescuing it to levels comparable to the untreated control. Data are represented as mean ± SD. Statistical significance is indicated by *p < 0.05, **p < 0.01, ***p < 0.001, and ****p < 0.0001 (ns = not significant).

### 3.2 Corilagin mitigates intracellular and mitochondrial oxidative stress

Given that hyperglycemia drives cellular damage primarily through oxidative stress, intracellular reactive oxygen species (ROS) were quantified using H2DCFDA staining. Flow cytometric histograms revealed a distinct rightward shift in DCF fluorescence in cells exposed to 50 mM glucose, indicating a robust generation of intracellular ROS. Pretreatment with 10 μM Corilagin significantly attenuated glucose-induced ROS generation, as evidenced by a leftward shift in fluorescence intensity and a significant reduction in MFI relative to the glucose-treated group (Figure 2A, B; p < 0.0001). To determine whether the increased ROS originated from mitochondria, mitochondrial superoxide production was assessed using MitoSOX™ Red staining. High glucose exposure significantly elevated mitochondrial superoxide levels compared with the control group (Figure 2C, D; p < 0.001). Corilagin pretreatment markedly suppressed mitochondrial ROS generation, significantly reducing MitoSOX™ fluorescence compared with glucose-treated cells (p < 0.0001), indicating that Corilagin effectively alleviates hyperglycemia-induced mitochondrial oxidative stress.

**Figure 2.**
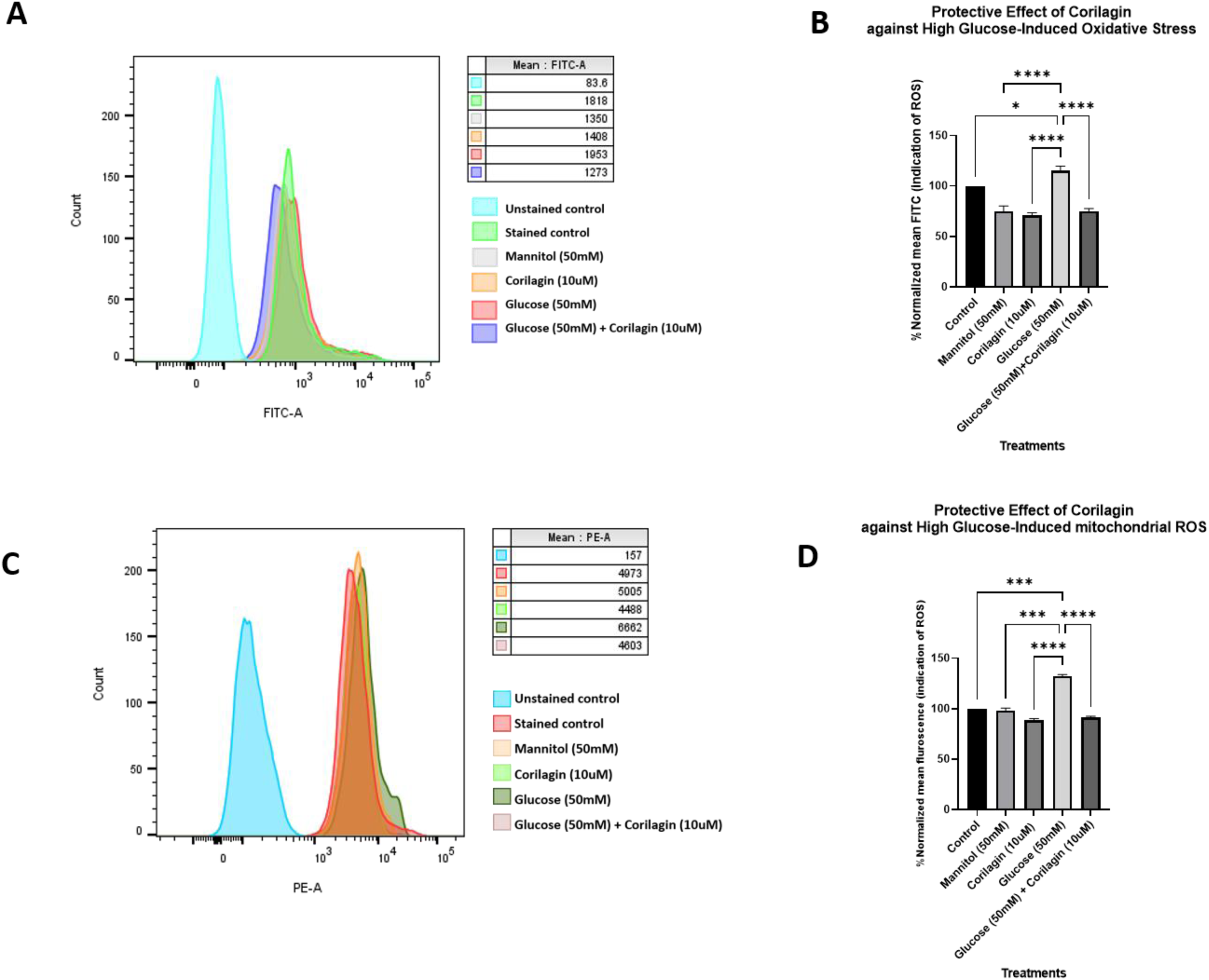
Corilagin attenuates glucose-induced intracellular and mitochondrial oxidative stress in SH-SY5Y cells. (A) Representative flow cytometric histograms of DCFH-DA (FITC) fluorescence showing intracellular reactive oxygen species (ROS) levels in SH-SY5Y cells following treatment with (1) control, (2) Corilagin (10 μM), (3) glucose (50 mM), and (4) Corilagin (10 μM) + glucose (50 mM). High-glucose exposure caused a marked increase in DCF fluorescence, indicative of elevated intracellular ROS generation, whereas co-treatment with Corilagin significantly reduced ROS accumulation toward basal levels. (B) Quantitative analysis of intracellular ROS expressed as mean fluorescence intensity (MFI), demonstrating a significant increase following glucose treatment and its attenuation by Corilagin. (C) Representative flow cytometric histograms of MitoSOX Red fluorescence showing mitochondrial superoxide production under the indicated treatment conditions. Glucose treatment markedly increased mitochondrial superoxide generation, while Corilagin co-treatment effectively suppressed mitochondrial oxidative stress. (D) Quantitative analysis of MitoSOX fluorescence (MFI) confirming a significant elevation of mitochondrial superoxide in glucose-treated cells and its significant reduction following Corilagin treatment. Data are presented as mean ± SD. Statistical significance is indicated by *P < 0.05, ****P < 0.0001

### 3.3 Corilagin preserves intracellular glutathione reserves

To assess the effect of Corilagin on the cellular antioxidant defense system, intracellular reduced thiol levels, primarily glutathione (GSH), were quantified using ThiolTracker™ staining. Exposure to 50 mM glucose significantly depleted intracellular reduced thiols, as evidenced by a leftward shift in the flow cytometric histogram and a significant reduction in MFI compared with the untreated control (Figure 3A, B; p < 0.05). Pretreatment with 10 μM Corilagin markedly restored intracellular reduced thiol levels, resulting in a significant increase in ThiolTracker™ fluorescence relative to the glucose-treated group (p < 0.0001).

**Figure 3.**
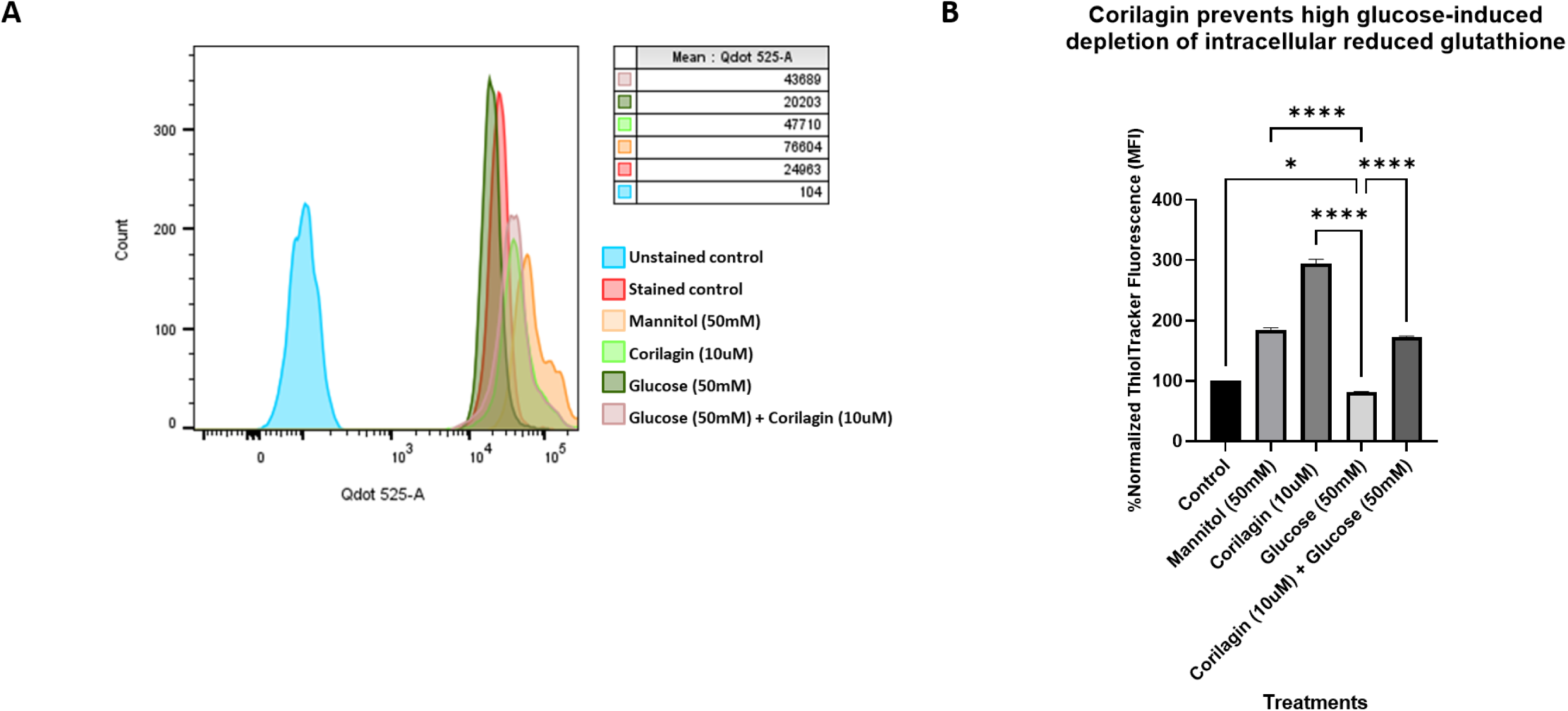
Corilagin protects against glucose-induced intracellular reduced glutathione depletion in SH-SY5Y cells. (A) Representative flow cytometric histograms of ThiolTracker Violet fluorescence showing intracellular reduced thiol levels in SH-SY5Y cells following treatment with (1) control, (2) Corilagin (10 μM), (3) glucose (50 mM), and (4) Corilagin (10 μM) + glucose (50 mM). Exposure to high glucose resulted in a marked leftward shift in fluorescence intensity, indicating depletion of intracellular GSH, whereas co-treatment with Corilagin restored ThiolTracker fluorescence toward control levels, suggesting preservation of the cellular antioxidant pool. (B) Quantitative analysis of ThiolTracker mean fluorescence intensity (MFI) demonstrating a significant reduction in intracellular reduced thiol content following glucose exposure and a significant recovery upon Corilagin treatment. Data are presented as mean ± SD. Statistical significance is indicated by *P < 0.05, ****P < 0.0001

### 3.4 Corilagin attenuates high glucose-induced cellular apoptosis

To investigate the protective effects of Corilagin against high glucose induced apoptosis, apoptotic cells were quantified following a 24-hour incubation period under various treatment conditions. Untreated control cells exhibited 7% apoptotic cells (Figure 4A). Treatment with 10 µM Corilagin alone did not significantly alter the apoptotic rate (∼8%), indicating that the compound exhibits no baseline cytotoxicity or pro-apoptotic effects at this dosage. Conversely, exposure to a hyperglycemic environment (50 mM glucose) induced a marked and statistically significant increase in cellular apoptosis, with the apoptotic cell population reaching approximately 16.5%. This elevation was highly significant when compared to both the untreated control group (*p* < 0.01) and the Corilagin-only control group (*p* < 0.01) (Figure 4B). Notably, co-treatment with 10 µM Corilagin and 50 mM glucose significantly attenuated the high glucose-induced apoptotic response, reducing the proportion of apoptotic cells to approximately 10.5%. This reduction was statistically significant (*p* < 0.05) relative to the cells treated with 50 mM glucose alone (Figure 4B).

**Figure 4.**
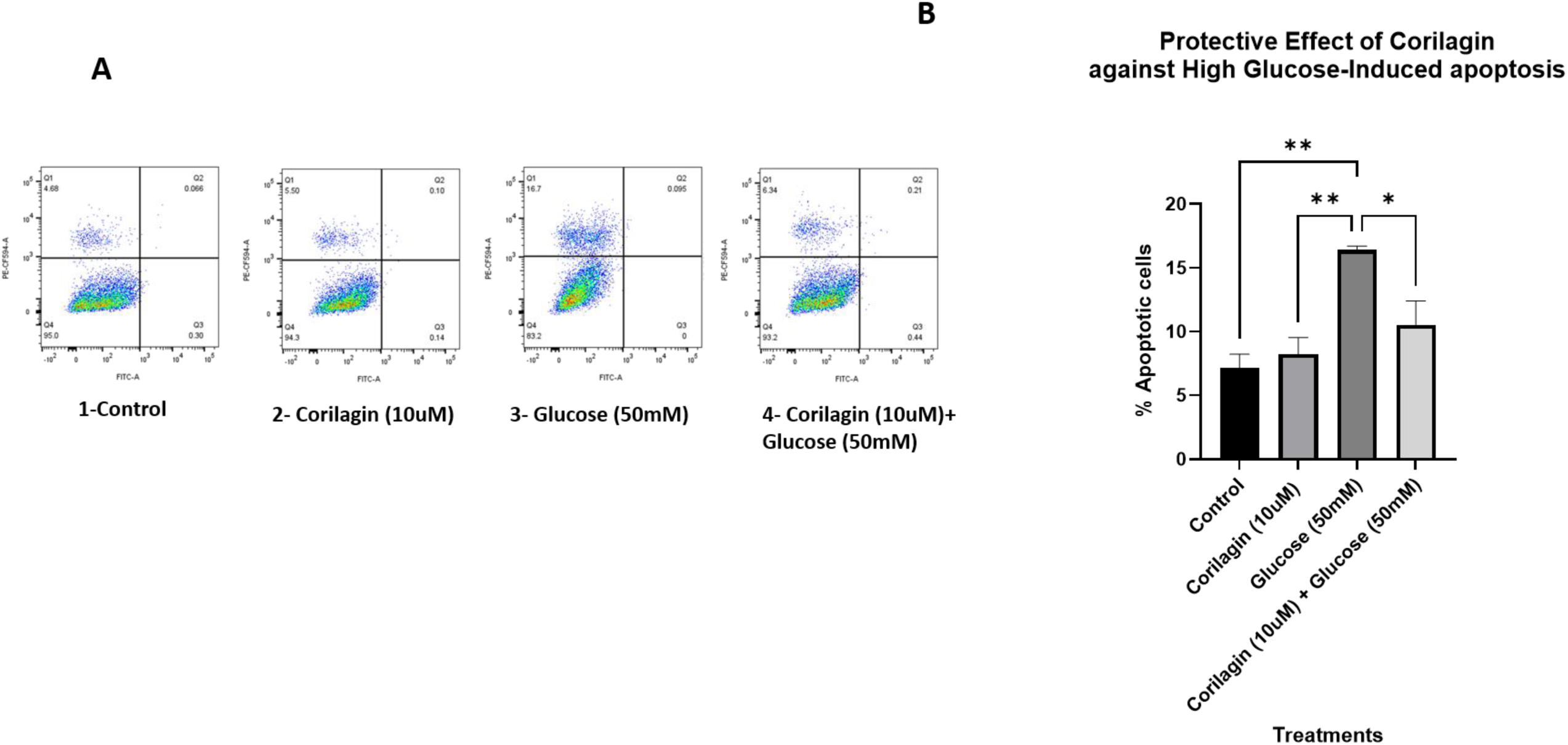
Corilagin attenuates glucose-induced apoptosis in SH-SY5Y cells. (A) Representative Annexin V-FITC/Propidium Iodide (PI) flow cytometric dot plots of SH-SY5Y cells following treatment with (1) control, (2) Corilagin (10 μM), (3) glucose (50 mM), and (4) Corilagin (10 μM) + glucose (50 mM). Cell populations were classified as viable (Annexin V⁻/PI⁻), early apoptotic (Annexin V⁺/PI⁻), late apoptotic (Annexin V⁺/PI⁺), and necrotic (Annexin V⁻/PI⁺). Exposure to high glucose markedly increased apoptotic cell populations, whereas co-treatment with Corilagin significantly reduced apoptosis and preserved cell viability. (B) Quantitative analysis of total apoptotic cells (early + late apoptosis) demonstrating a significant increase following glucose treatment, which was substantially reversed by Corilagin treatment. Data are presented as mean ± SD. Statistical significance is indicated by *P < 0.05, ** P < 0.01

### 3.5 Corilagin maintains mitochondrial membrane potential

The preservation of mitochondrial function was further evaluated by measuring the mitochondrial membrane potential (Δψ_m_) using JC-1 staining. Confocal microscopy revealed that healthy control cells exhibited strong red fluorescence (J-aggregates), indicative of a high, polarized membrane potential. Following exposure to 50 mM glucose, there was a pronounced loss of red fluorescence and a concomitant increase in green fluorescence (monomers), signifying mitochondrial depolarization and dysfunction. Quantitative analysis revealed a significant reduction in the red-to-green fluorescence ratio compared with the untreated control (p < 0.0001; Figure 5B). Pretreatment with 10 μM Corilagin markedly preserved mitochondrial membrane potential, as evidenced by restoration of red fluorescence and a significantly higher red-to-green fluorescence ratio compared with the glucose-treated group (p < 0.0001), approaching that observed in the untreated control.10µM Corilagin pretreatment successfully preserved mitochondrial polarity, exhibiting a high red-to-green fluorescence ratio that closely resembled the untreated control group (Figure 5A, B).

**Figure 5.**
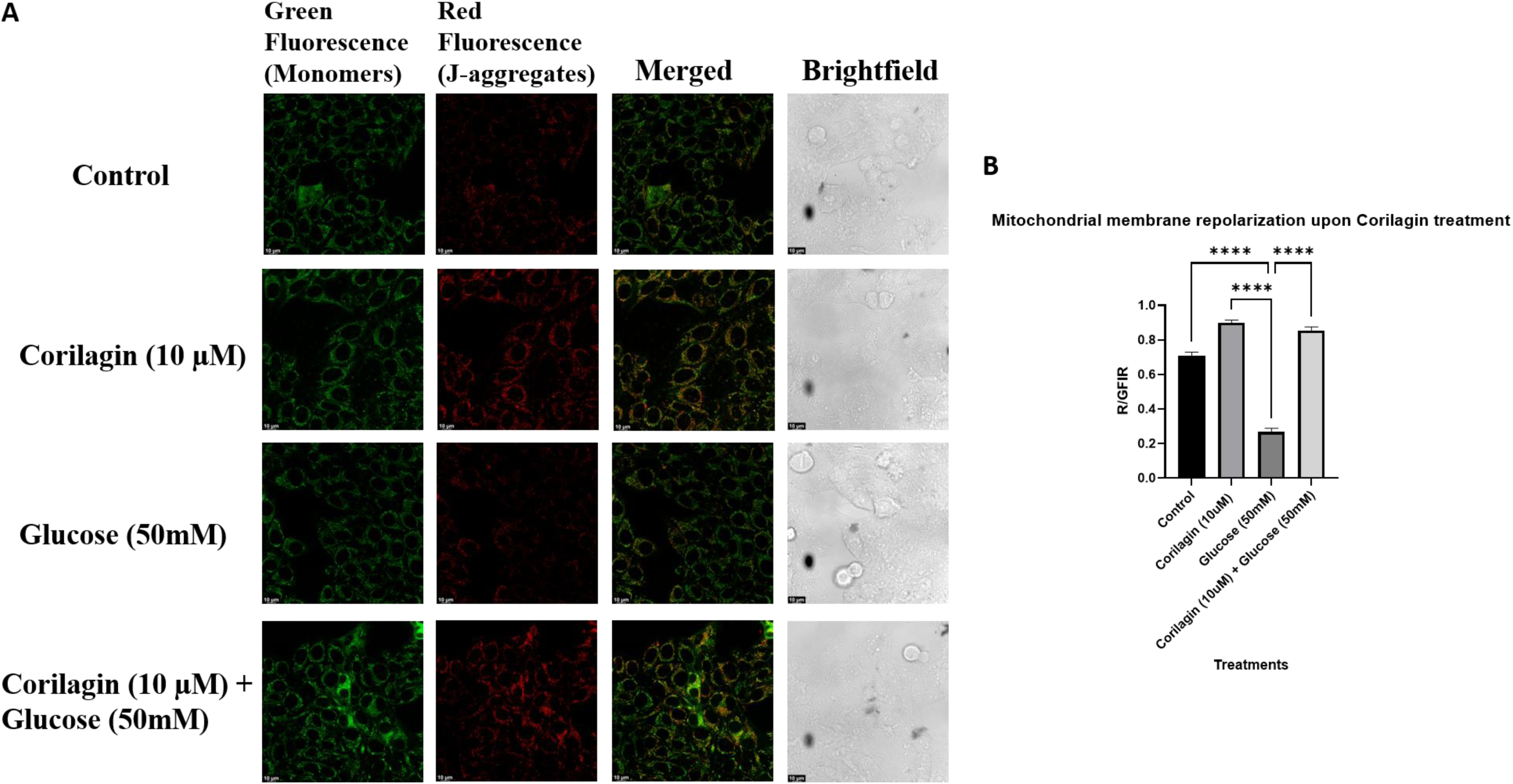
Corilagin preserves mitochondrial membrane potential against high glucose-induced depolarization. **(A)** Representative confocal fluorescence and differential interference contrast (DIC) microscopy images of cells stained with the mitochondrial polarity indicator JC-1. In healthy cells (Control and Corilagin alone), JC-1 accumulates in the mitochondria as J-aggregates, emitting strong red fluorescence indicative of a high Δψ_m_. Exposure to 50 mM glucose caused a marked reduction in red fluorescence and an increase in green fluorescence (JC-1 monomers), indicating severe mitochondrial depolarization and dysfunction. Pretreatment with 10 µM Corilagin effectively preserved mitochondrial structural integrity and membrane polarity, restoring the red J-aggregate fluorescence. Scale bars = 10 µm. **(B)** Quantitative analysis of the mitochondrial membrane potential, represented by the ratio of red to green fluorescence intensity. The 50 mM glucose challenge resulted in a significant decrease in the red/green ratio compared to the untreated control. Corilagin pretreatment successfully rescued the membrane potential, returning the red/green ratio to near-baseline levels. Data are presented as mean ± SD. Statistical significance is indicated by ****p < 0.0001

### 3.6 Corilagin attenuates high glucose-induced mitochondrial fragmentation

To evaluate the protective effects of Corilagin on mitochondrial dynamics under hyperglycemic stress, mitochondrial morphology was assessed using MitoTracker staining and subsequently quantified (Figure 6A). Four experimental groups were analysed for structural parameters such as roundness, area, fibre aspect ratio and Feret maximum; Control, Corilagin (10 µM), Glucose (50 mM) and co-treatment of Corilagin (10 µM) + Glucose (50 mM). Quantitative analysis showed that mitochondrial fragmentation was severe under high glucose (50 mM) exposure. Mitochondria in glucose treated cells showed a significant increase in roundness compared to the untreated control (p<0.01), indicating a change from elongated networks to fragmented, punctate structures. Marked decreases in general mitochondrial area (p < 0.001), fibre aspect ratio (p < 0.05) and Feret maximum (p < 0.001) also further support this fragmentation confirming a loss of tubular mitochondrial networks. However, co-treatment with 10 µM Corilagin strongly rescued the mitochondrial network from the glucose-induced damage. Corilagin addition to the high glucose environment strongly reversed mitochondrial rounding (p<0.0001) (Figure 6B. Furthermore, it significantly ameliorated and restored mitochondrial elongation and size, as evidenced by highly significant increases in mitochondrial area (p < 0.0001), fibre aspect ratio (p < 0.0001) and Feret maximum (p < 0.0001) when compared to the glucose-only group. In particular, the co-treatment group showed higher levels of these structural parameters than baseline controls, suggesting a significant increase in mitochondrial fusion. Moreover, in cells treated with Corilagin alone (10 µM) there was a clear tendency of increased mitochondrial area and fibre aspect ratio in comparison with the control, suggesting that Corilagin per se promotes a fused, elongated mitochondrial phenotype in the absence of metabolic stress (Figure 6B).

**Figure 6.**
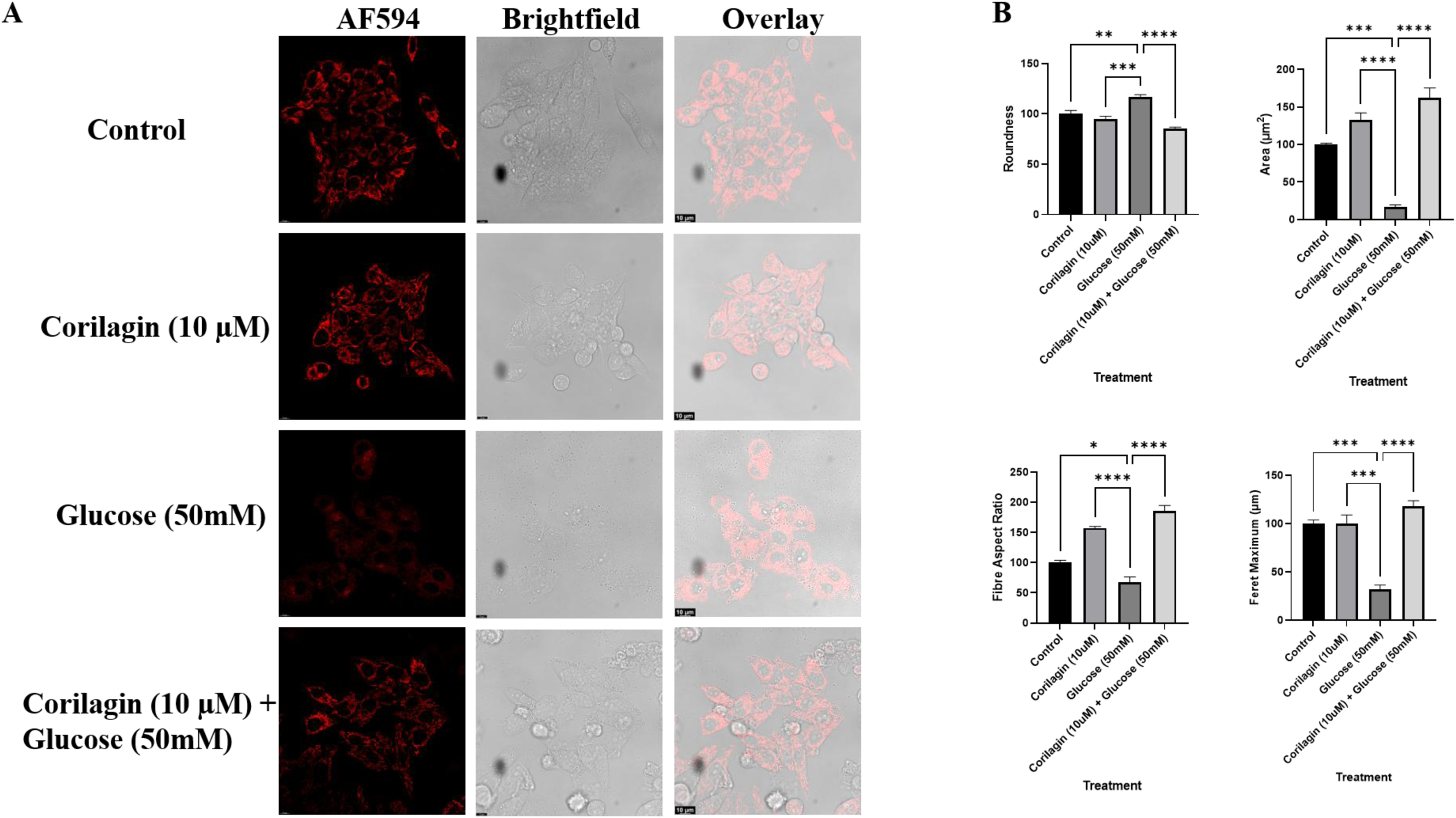
Corilagin preserves mitochondrial morphology in SH-SY5Y cells under hyperglycaemic conditions. (A) Representative confocal microscopy images of SH-SY5Y cells stained with MitoTracker showing mitochondrial morphology under four experimental conditions: (1) control, (2) Corilagin (10 μM), (3) glucose (50 mM), and (4) Corilagin (10 μM) + glucose (50 mM). Brightfield and merged images are shown alongside the fluorescence images. Scale bar = 10 μm. (B) Quantitative morphometric analysis of mitochondrial morphology, including roundness, mitochondrial area, fibre aspect ratio, and Feret maximum, demonstrating significant mitochondrial fragmentation following glucose exposure and its attenuation by Corilagin treatment. Data are presented as mean ± SD. Statistical significance is indicated by *P < 0.05, **P < 0.01, ***P < 0.001, ****P < 0.0001.

### 3.8 *Corilagin confers mitochondrial protection via the AMPK-SIRT1-PGC1*α *signaling axis*

To elucidate the molecular mechanisms underlying the cytoprotective and mitochondria-preserving effects of Corilagin, the expression of key proteins involved in mitochondrial biogenesis was analyzed by Western blotting. Exposure to 50 mM glucose significantly reduced the phosphorylation of AMPK (P-AMPK) without affecting total AMPK expression (Figure 7A, B; p < 0.05). This was accompanied by a significant downregulation of the downstream regulators SIRT-1, PGC1α and TFAM compared with the untreated control (p < 0.01, p < 0.001). Pretreatment with 10 μM Corilagin significantly restored P-AMPK levels and markedly increased the expression of SIRT-1, PGC1α, and TFAM relative to the glucose-treated group (p < 0.05, p < 0.001), with protein levels approaching those observed in the control group. These findings indicate that Corilagin alleviates hyperglycemia-induced mitochondrial dysfunction through activation of the AMPK–SIRT1–PGC1α–TFAM signaling axis.

**Figure 7.**
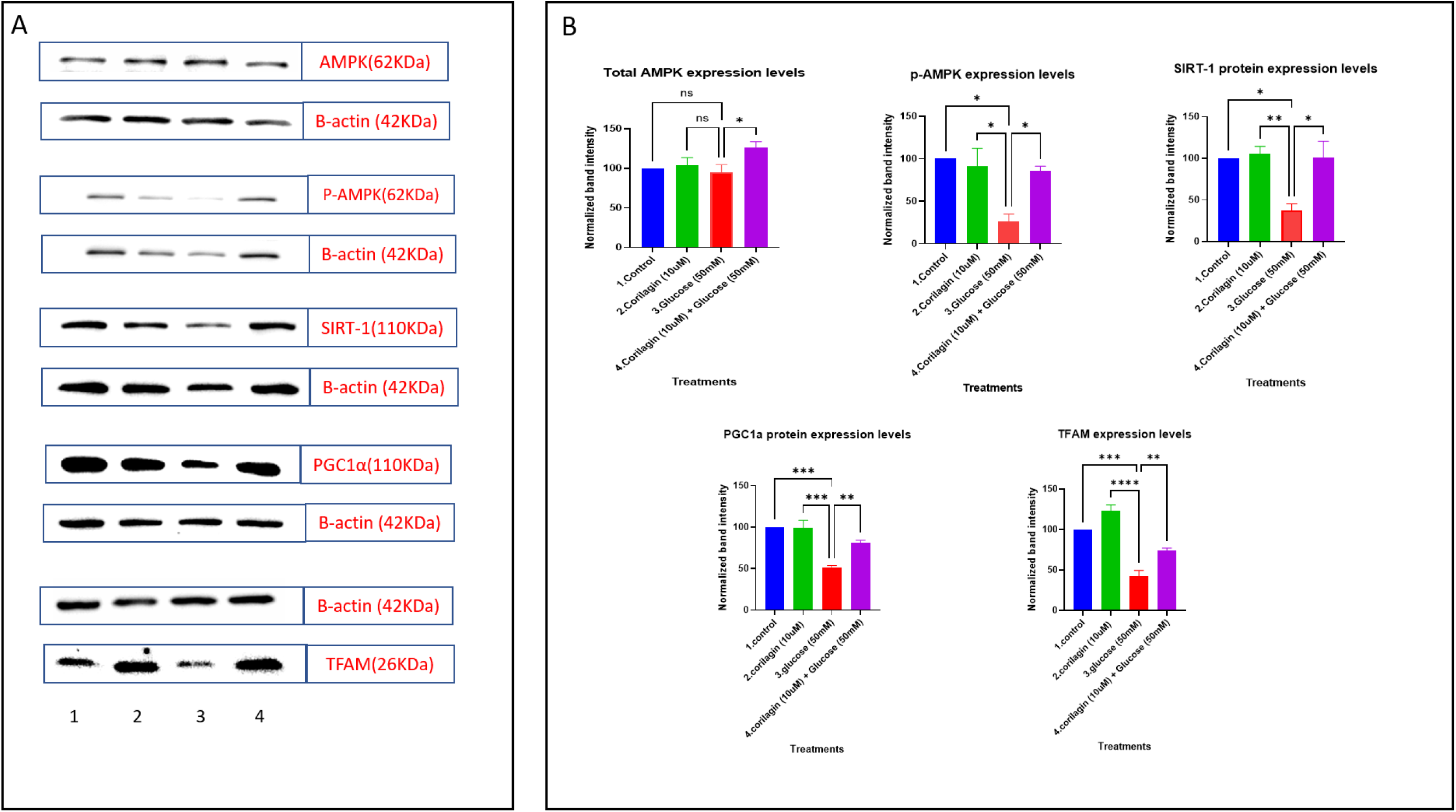
Corilagin activates the AMPK-SIRT1-PGC-1*α* signaling axis to mitigate high glucose-induced cellular stress. (A) Representative Western blot images depicting the protein expression profiles of total AMPK, phosphorylated AMPK (P-AMPK), SIRT-1, PGC-1α, and TFAM in SH-SY5Y cells. β-actin was utilized as an endogenous loading control to ensure equal protein concentration across lanes. The numbered lanes correspond to the following experimental groups: 1 (Untreated Control), 2 (10 µM Corilagin alone), 3 (50 mM Glucose alone), and 4 (10 µM Corilagin pretreatment + 50 mM Glucose). (B) Densitometric quantification of the Western blot bands, presented as normalized band intensity. Exposure to 50 mM glucose significantly suppressed the phosphorylation of AMPK (P-AMPK) and markedly downregulated the expression of downstream metabolic and mitochondrial biogenesis regulators, including SIRT-1, PGC-1α, and TFAM. Pretreatment with 10 µM Corilagin successfully rescued this suppression, significantly upregulating the expression of these critical protective proteins. Total AMPK expression levels remained largely stable across the treatment groups, indicating that Corilagin specifically modulates the functional activation (phosphorylation) of AMPK rather than its basal translation. Data are expressed as mean ± SD. Statistical significance between indicated groups is denoted by brackets: *p < 0.05, p < 0.01, ***p < 0.001, and ****p < 0.0001 (ns = not significant).

## 4. Discussion

Mitochondrial dysfunction is one of the central biochemical mechanisms that drives hyperglycemia-induced neuronal injury. In diabetic neuropathy models, the excessive glucose flux into neurons causes mitochondrial superoxides to accumulate in high quantities. The subsequent increase in the cellular level of superoxides leads to an overproduction of reactive oxygen species (ROS), initiating and driving the apoptotic process within the neuron [4, 27, 28]. Using SH-SY5Y cells as an in-vitro model for studying the effects of hyperglycemic shock on neurons, we demonstrated that Corilagin treatment significantly protected both the mitochondrial structure and cell viability against the damaging effects of high glucose concentrations. The protective effects of Corilagin were coincident with the reactivation of the AMPK–SIRT1–PGC1α–TFAM signaling pathway signaling axis. A significant elevation in both total cellular and specifically mitochondrial ROS occurred early in the sequence of events leading to neuronal damage in our model. Furthermore, there was a corresponding decrease in reduced glutathione levels. This pattern of events is consistent with Brownlee’s unifying model of hyperglycemic mitochondrial overload. According to this model, excessive glucose flux into neurons results in the generation of high amounts of superoxide radicals at the site of the mitochondria. These radicals then initiate all of the major pathogenetic mechanisms responsible for diabetes-related complications [4, 29]. In addition to blocking the formation of ROS, Corilagin may enhance endogenous antioxidant defenses through activation of the Nrf2 signaling pathway. Nrf2 regulates the transcription of numerous antioxidant and glutathione-metabolizing enzymes, including glutamate-cysteine ligase, glutathione reductase, heme oxygenase-1, and NADPH quinone oxidoreductase-1, thereby preserving intracellular glutathione homeostasis under oxidative stress. Notably, Corilagin has previously been shown to activate Nrf2, restore glutathione levels, increase antioxidant enzyme activity, and reduce oxidative damage in experimental cerebral ischemia, suggesting that a similar mechanism may contribute to the preservation of intracellular thiols observed in the present study [21, 30]. Following this initial oxidative stress, high glucose led to an extensive depolarization and disintegration of mitochondrial networks. Depolarization and fragmentation of mitochondrial networks are consistent with previous observations demonstrating a direct relationship between oxidative stress and dysfunctional mitochondrial quality control in various animal models of diabetic neuropathy [17, 18].

Treatment with Corilagin prior to exposure to high glucose prevented the depolarization and fragmentation observed when exposed to high glucose alone. Several morphometric measurements taken from cells treated with Corilagin + high glucose revealed greater numbers of intact mitochondrial networks than cells treated with high glucose alone. This suggests that Corilagin not only prevents the detrimental effects of high glucose on mitochondrial function but may also enhance mitochondrial fusion. However, because enhanced fusion does not necessarily indicate an improvement in mitochondrial health, caution should be exercised when interpreting these findings. Enhanced fusion can represent either an adaptative response to stress or a compensatory response to damaged mitochondria. Further research using measures of mitochondrial respiratory function will be necessary to determine if the increased fusion observed in Corilagin-treated cells represents true biogenic enhancement or merely a stressful morphological adaptation. The excess ROS production and mitochondrial membrane depolarization trigger apoptosis in neural cell undergone with hyper-glycemic shock. Pretreatment with Corilagin rescues cells from apoptosis by dampening oxidative burden.

At the molecular level, high glucose suppressed AMPK phosphorylation and decreased SIRT1, PGC1α, and TFAM protein abundance. Conversely, Corilagin restored each of these changes. AMPK, SIRT1, and PGC1α are members of a highly interconnected energy-sensing regulatory network. Activation of AMPK increases NAD+ availability, which activates SIRT1, which subsequently deacetylates and activates PGC1α to regulate mitochondrial biogenesis via TFAM. Activation of each member reinforces the cycle [31].

This pattern of activation is consistent with previous studies demonstrating that Corilagin activates the SIRT1-AMPK pathway in diabetic nephropathy models to improve podocyte autophagy and reduce oxidative stress caused by high glucose [20]. Similar pattern of activation has been observed in Schwann cells derived from diabetic peripheral neuropathy models treated with another natural compound (honokiol). In these models, honokiol activated the same AMPK-SIRT1-PGC1α axis to protect Schwann cells from high glucose injury. Furthermore, resveratrol has been shown to activate AMPK/SIRT1 signaling, restore PGC1α and TFAM expression, improve mitochondrial biogenesis, reduce oxidative stress and improve nerve function in experimental diabetic neuropathy. Likewise, quercetin and berberine, have all demonstrated neuroprotective effects by enhancing AMPK–SIRT1–PGC1α signaling, preserving mitochondrial integrity and reducing ROS generation in Streptozotocin (STZ) induced diabetic rat models [32–35]. Collectively, these findings support the concept that restoration of mitochondrial bioenergetics through the AMPK–SIRT1–PGC1α axis represents a convergent mechanism underlying the neuroprotective actions of several plant-derived compounds, with our findings identifying Corilagin as another promising modulator of this pathway. Collectively, these findings suggest that the AMPK-SIRT1-PGC1α-TFAM signaling axis may be a common pathway through which Corilagin exerts protective effects across different tissues.

However, our findings demonstrate only a correlation between AMPK phosphorylation-SIRT1-PGC1α-TFAM levels and the protective phenotypes observed in Corilagin-treated cells. Future pharmacological studies employing inhibitors of AMPK or knockdown of overexpressed genes or transcripts will be essential for establishing whether this pathway is mechanistically required for Corilagin’s cytoprotective actions or if it plays one among multiple roles contributing to overall protection afforded by Corilagin. Mitochondrial ATP production/OXPHOS capacity was not directly measured, so it remains unclear whether Corilagin’s protective effects extend to genuine bioenergetic restoration beyond the structural and redox improvements observed. Overall, Corilagin reduced oxidative stress, mitochondrial dysfunction and apoptosis induced by high glucose concentration in SH-SY5Y cells; these reductions were associated with activation of AMPK-SIRT1-PGC1α-TFAM signaling. Based upon these findings, Corilagin appears to act as a novel mitochondrially-targeted neuroprotective agent warranting further evaluation as such using primary neuronal cultures and in vivo models of diabetic neuropathy.

## List of Abbreviations

AGEs: Advanced Glycation End Products
AMPK: AMP-Activated Protein Kinase
ANOVA: Analysis of Variance
AR: Aspect Ratio
ATP: Adenosine Triphosphate
BCA: Bicinchoninic Acid
BSA: Bovine Serum Albumin
COß: Carbon Dioxide
DCF: 2′,7′-Dichlorofluorescein
DCFDA (HßDCFDA): 2′,7′-Dichlorodihydrofluorescein
Diacetate DMEM: Dulbecco’s Modified Eagle Medium
DMSO: Dimethyl Sulfoxide
DN: Diabetic Neuropathy
DPBS: Dulbecco’s Phosphate-Buffered Saline
ECL: Enhanced Chemiluminescence
FBS: Foetal Bovine Serum
FITC: Fluorescein Isothiocyanate
GSH: Reduced Glutathione
HRP: Horseradish Peroxidase
JC-1: 5,5′,6,6′-Tetrachloro-1,1′,3,3′-tetraethylbenzimidazolylcarbocyanine iodide
MFI: Mean Fluorescence Intensity
MTT: 3-(4,5-Dimethylthiazol-2-yl)-2,5-diphenyltetrazolium Bromide
Nrf2: Nuclear Factor Erythroid 2-Related Factor 2
OXPHOS: Oxidative Phosphorylation
P-AMPK: Phosphorylated AMP-Activated Protein Kinase
PBS: Phosphate-Buffered Saline
PE: Phycoerythrin
PGC1α: Peroxisome Proliferator-Activated Receptor Gamma Coactivator-1 Α
PI: Propidium Iodide
PVDF: Polyvinylidene Difluoride
RIPA: Radioimmunoprecipitation Assay
ROS: Reactive Oxygen Species
SDS-PAGE: Sodium Dodecyl Sulfate–Polyacrylamide Gel Electrophoresis
SIRT1: Sirtuin 1
TBST: Tris-Buffered Saline with Tween-20
TFAM: Mitochondrial Transcription Factor A
Δψm: Mitochondrial Membrane Potential

## Author contribution

RC- Conceptualization, Investigations, Methodology, Data Curation, Analysis, Writing Original Draft.

DR- Investigations, Methodology, Data Curation, Analysis, Writing Original draft.

JC- Investigations, Methodology, Data Curation, Analysis

BB- Investigations, Data Curation, Analysis.

MA-Investigations, Data Curation, Analysis.

MM- Investigations, Data Curation, Analysis.

DG- Conceptualization, Analysis, Writing- Review and Editing, Supervision, Resources.

## Acknowledgement and declaration (justified use of AI tools)

All the authors would like to acknowledge National Institute of Pharmaceutical Education and Research (NIPER) Kolkata, Ministry of Chemicals and Fertilisers, Government of India, for providing infrastructure and fundings to carry out the research work. Author RC would like to acknowledge DST INSPIRE, Govt. of India. DR acknowledges UGC, Govt. of India. Authors JC and BB, would like to acknowledge the Department of Pharmaceuticals, Ministry of Chemicals and Fertilisers, Govt. of India; Author MA would like to acknowledge ICMR, Govt. of India for providing fellowship. Authors would like to declare the usage of Google Gemini Pro for the improvement of the English language. After using this tool/service, the author(s) reviewed and edited the content as needed and take(s) full responsibility for the content of the published article.

## Funding Declaration

The authors declare no funds were received during the preparation of manuscript.

## Conflict of Interest

The authors declare that they have no conflict of interest.

## Notes

### Competing Interest Statement

The authors have declared no competing interest.

